# Analysis of β rhythm induction in acute brain slices using field potential imaging with ultra-high-density CMOS-based microelectrode array

**DOI:** 10.1101/2025.03.30.646248

**Authors:** Hiroto Takahashi, Naoki Matsuda, Ikuro Suzuki

## Abstract

Carbachol-induced beta rhythms in acute brain slices provide a promising ex vivo platform for evaluating compound effects under arousal-like hippocampal activity. Although microelectrode array (MEA) systems offer a non-invasive means of recording network dynamics, conventional MEAs have been limited by low electrode counts and small sensing areas, allowing only fragmented evaluation of neural activity. In this study, we applied field potential imaging (FPI) using an ultra-high-density (UHD) CMOS-MEA comprising 236,880 electrodes to simultaneously and spatially resolve carbachol-induced activity changes in the hippocampus and neocortex. Treatment with 10 µM carbachol reliably induced oscillatory activity in the hippocampus at 1.33 ± 0.33 Hz. Repetitive and temporally precise firing patterns originated in the CA3 region and propagated toward CA1. The average duration of individual oscillatory events was 52.82 ± 6.44 ms, corresponding to a frequency of 18.93 ± 3.03 Hz—within the beta frequency band. Cluster analysis based on waveform morphology revealed spatial groupings that reflected underlying anatomical structures. Beta-band power was significantly enhanced in both hippocampal and cortical regions following carbachol treatment. These findings demonstrate that the FPI platform enables high-resolution mapping of relationships between anatomical structures and brain function in acute slices, and holds promise as a pharmacological evaluation tool for assessing compound effects on hippocampal and cortical networks under wakefulness-like conditions.

## Introduction

Beta rhythms observed in the hippocampus are electrical oscillations typically associated with wakeful states. These oscillations are thought to play a central role in attentional control, problem-solving, creative cognition, and sensory information processing during arousal (Kay et al., 2009;Engel and Fries, 2010). They are closely linked to specific neural circuits, including cholinergic projections from the medial septum to the hippocampal CA3 region (Shimono et al., 2000;Teles-Grilo Ruivo and Mellor, 2013). Dysregulation of beta activity has been implicated in neurological disorders such as Parkinson’s disease and Alzheimer’s disease (Hammond et al., 2007;Engel and Fries, 2010;Little and Brown, 2014).

Thus, investigating beta rhythms not only deepens our understanding of the electrophysiological basis of cognitive function but also provides critical insights for elucidating disease mechanisms and developing therapeutic strategies. Capturing the complex dynamics of neural activity in real time is essential for decoding brain function, including the generation and propagation of beta oscillations. In vivo recordings using microelectrodes enable precise targeting of specific brain regions and have been instrumental in revealing both single-neuron and population-level activity (Le Van Quyen et al., 2010). However, despite their widespread use in neuroscience and disease modeling, in vivo techniques have intrinsic limitations, including their invasiveness, limited spatial coverage, and reduced throughput for compound screening.

By contrast, ex vivo approaches, such as acute brain slice preparations, offer high-resolution access to individual neurons and microcircuits, making them particularly suitable for analyzing the electrophysiological and pharmacological properties of local neural networks. Among these, microelectrode array (MEA) technology has become a valuable tool for noninvasive, simultaneous recordings from hippocampal subfields—namely CA1, CA3, and the dentate gyrus (DG) (Keefer et al., 2008). Nevertheless, conventional MEAs typically consist of only 16 to 64 electrodes, limiting their ability to resolve circuit-level dynamics or capture activity from broader brain regions beyond the hippocampus.

Recent advances in complementary metal-oxide-semiconductor (CMOS)-based MEAs have overcome many of these limitations by offering ultra-dense electrode integration with high spatial and temporal resolution (Frey et al., 2010;Ballini et al., 2014). We previously developed a field potential imaging (FPI) platform using an ultra-high-density (UHD) CMOS-MEA composed of 236,880 electrodes spanning a 5.5 × 5.9 mm sensing area. This system enables whole-slice recordings across hippocampal and cortical regions with unprecedented resolution (Suzuki et al., 2023). Despite these technical advances, beta oscillations do not spontaneously occur in brain slices under standard ex vivo conditions, in contrast to in vivo. For pharmacological studies to model awake-state activity more effectively, it is essential to establish an induced brain state that replicates arousal-related electrophysiology. If beta rhythms can be reliably induced, in vitro preparations may provide a valuable framework for evaluating compound effects under wakefulness-like conditions.

In this study, we used the UHD-CMOS-MEA platform to record hippocampal and cortical activity during carbachol-induced beta rhythm. We simultaneously monitored multiple brain regions, characterized the evoked oscillatory dynamics, and examined region-specific differences in beta-band power within hippocampal subfields and associated cortical areas.

## Materials and Methods

### Mouse acute brain Slice

Acute brain slice experiments were conducted using 6- to 7-week-old male mice (C57BL/6NCrSlc, Japan SLC, Inc.). The mice were decapitated under isoflurane (Viatris Inc.) inhalation anesthesia, and the brain was extracted. The cerebellum and the anterior 1/3 of the cerebrum were removed. The brain was placed with the rostral side facing down, and approximately 20–30 degrees of the left and right hemispheres were trimmed. The trimmed brain was divided into right and left hemispheres, positioned dorsal side up, and sliced into 300 µm-thick sections using a slicer (Neo-LinearSlicer NLS-MT, DOSAKA EM CO., LTD.). The experiments were conducted following the animal experiment guidelines approved by Tohoku Institute of Technology (Approval Number: 2020-01). The brain slices were incubated for 1 hour at 30–32°C for recovery and maintained at room temperature until measurement. The recovery solution consisted of artificial cerebrospinal fluid (aCSF) containing 124 mM NaCl, 3.0 mM KCl, 26 mM NaHCO□, 1.25 mM KH□PO□, 2.4 mM CaCl□·2H□O, 1 mM MgSO□·7H□O, and 10 mM D-(+)-glucose, and was continuously bubbled with a 95% O□/ 5% CO□gas mixture.

### Ultra-high-density (UHD)-CMOS-MEA measurement system

The UHD-CMOS-MEA (Sony Semiconductor Solutions, Co., Ltd.) features a total of 236,880 electrodes, each measuring 10.52 µm × 10.52 µm, arranged at an interval of 1.20 µm within a sensing area of 5.9 mm × 5.5 mm. These electrodes are organized in a 504-row × 470-column matrix (Suzuki et al., 2023). The MEA chip was mounted onto a dedicated MEA unit, and the signals acquired from all electrodes were transmitted as binary data to a PC via an FPGA unit (Figure 1A). During spontaneous activity recording, a perfusion cap was placed over the CMOS-MEA chip, and low-magnesium (0.1 mM) aCSF was circulated at 20 rpm using a peristaltic pump (Gilson: 61-0083-52, Mini Pulse Pump III MP-4 F155006). All 236,880 electrodes were simultaneously activated and recorded at a sampling rate of 1 kHz. The recorded waveform data were visualized using Bin Viewer CMOS software (SCREEN, Co., Ltd.).

**Figure 1.**
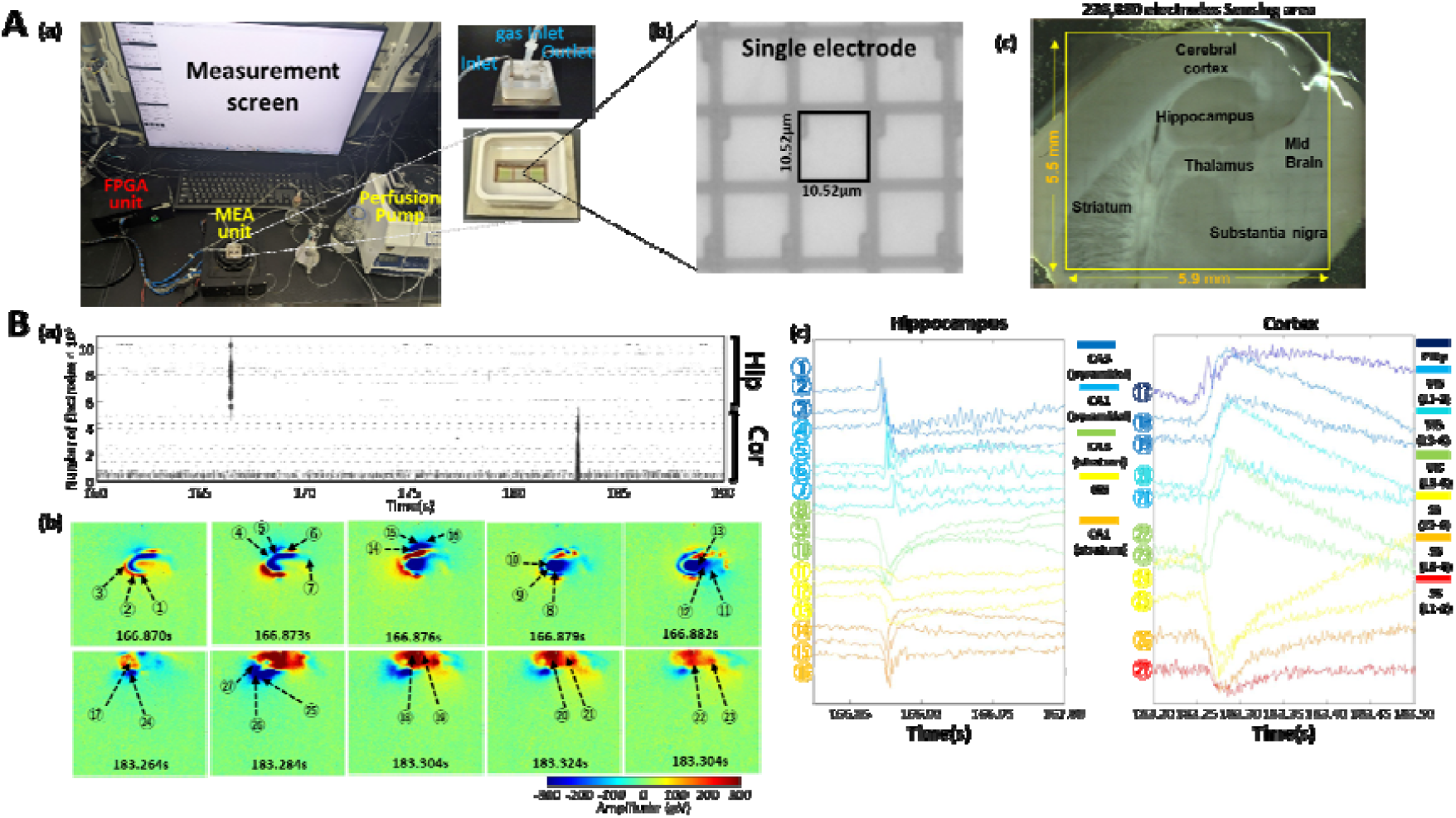
Acute Brain Slice Recording Using an Ultra-high-density (UHD) CMOS-MEA. (A) (a) Experimental setup for acute brain slice recording using the UHD-CMOS-MEA, including the perfusion system and FPGA interface. (b) Optical microscope image showing a single electrode on the UHD-CMOS-MEA (10.52 µm × 10.52 µm). (c) Acute sagittal mouse brain slice mounted on the UHD-CMOS-MEA, with major anatomical regions labeled within the 5.5 mm × 5.9 mm sensing area. (B) (a) Raster plot of spontaneous activity recorded under perfusion with low-magnesium aCSF, highlighting time points of activation in the hippocampal and cortical regions. (b) Spatial potential maps showing hippocampal and cortical region-specific activations at different time points. Red and blue indicate positive and negative potentials, respectively. (c) Raw extracellular waveforms corresponding to activation events shown in (b). Waveform numbers correspond to the regions indicated in panel B(b).

### Pharmacological Testing

Acute mouse brain slices were mounted onto the UHD-CMOS-MEA, followed by a 15-minute stabilization period. After stabilization, spontaneous neural activity was recorded for 5 minutes. Subsequently, to induce beta rhythm, carbachol (10 µM, Wako) was applied via perfusion. Ten minutes after the onset of perfusion, neural activity was recorded again for 5 minutes.

### Signal Detection

The acquired extracellular voltage waveforms were processed using a bandpass finite impulse response (FIR) filter ranging from 1 to 100□Hz. Neural activity signals were then detected based on a voltage threshold method. For visualization of the detected events, threshold crossing times were plotted only for electrodes exhibiting high signal amplitudes, using a detection threshold of ±200 μV. The baseline noise level was 22.08□±□5.28 μV, and the 200 μV threshold corresponds to approximately 9 times the standard deviation. Signal detection and visualization were performed using MATLAB (MathWorks, Natick, MA).

### Cluster Classification

From the total of 236,880 electrodes, a subset of 155,365 active electrodes was extracted using a voltage threshold of ±80 μV. K-means clustering was then performed on the 5-second extracellular voltage waveforms recorded from these active electrodes. The optimal number of clusters was determined using the elbow method. Based on the clustering results, a cluster heatmap was generated by assigning colors to the electrodes according to their cluster membership. All clustering procedures and visualizations were performed using MATLAB (MathWorks, Natick, MA).

### Frequency Analysis

Frequency analysis was conducted for each cluster-defined region. First, the raw voltage waveforms from all electrodes within each cluster were transformed using fast Fourier transform (FFT) to obtain individual frequency spectra. FFT was performed on 5-second recordings (5000 samples). These spectra were then averaged within each cluster to generate cluster-specific spectral profiles. Next, using the averaged frequency spectra at 0.2 Hz resolution, cluster-based frequency heatmaps were generated (Figure 3B). Additionally, the mean spectral power was calculated for the following frequency bands: delta (1–3 Hz), theta (4–8 Hz), alpha (8–15 Hz), beta (15–30 Hz), gamma (35–50 Hz), high-gamma (80–150 Hz), and ripple (150–200 Hz). All FFT computations and heatmap visualizations were performed using MATLAB (MathWorks, Natick, MA).

**Figure 2.**
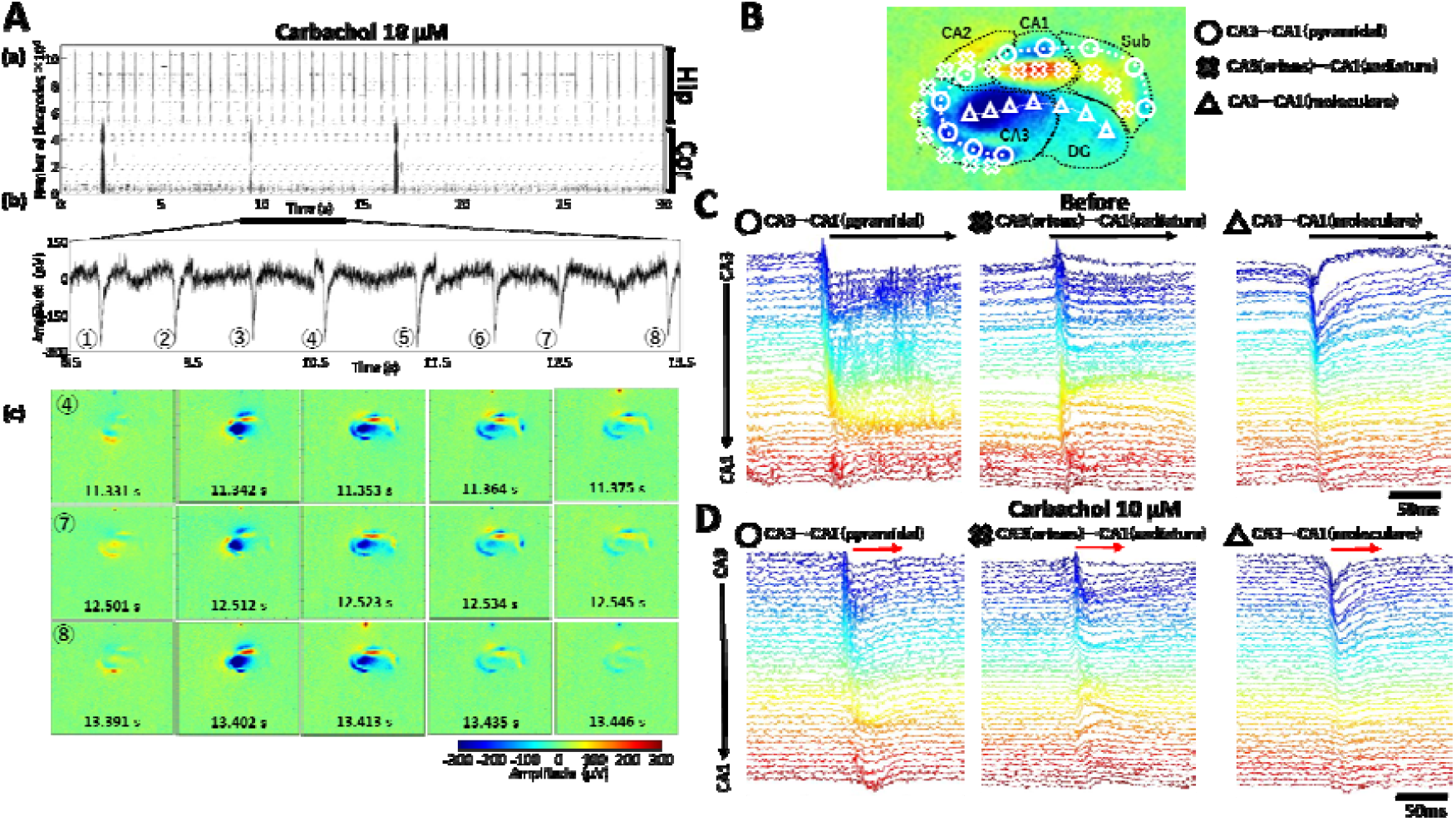
Induction of Beta Rhythm in the Hippocampus by Carbachol Treatment. (A) (a) Raster plot of hippocampal and cortical activity recorded over 30 seconds following carbachol treatment under low-magnesium aCSF perfusion. (b) Raw extracellular waveform recorded from a single electrode in the hippocampal region, showing repetitive oscillatory activity. (c) Time-series potential maps of hippocampal activity corresponding to oscillatory events ➃, ➆, and ➇ from panel A(b). Each row represents consecutive time frames (left to right) during a single oscillatory event. (B) Maximum amplitude potential map during carbachol-induced oscillation, with overlaid arrows indicating propagation directions. Anatomical landmarks such as CA2, CA1, CA3, DG, and Subiculum are labeled. The three propagation pathways from CA3 to CA1 are marked : pyramidal layer (○), oriens–radiatum (×), and molecular–molecular (△). (C) Raw waveforms representing activity propagation along each identified pathway during carbachol treatment.Left: From CA3 pyramidal layer to CA1 pyramidal layer (○). Center: From CA3 oriens layer to CA1 radiatum (×). Right: From CA3 molecular layer to CA1 molecular layer (△). Waveforms are color-coded according to electrode position from dorsal (red) to ventral (blue).

**Figure 3.**
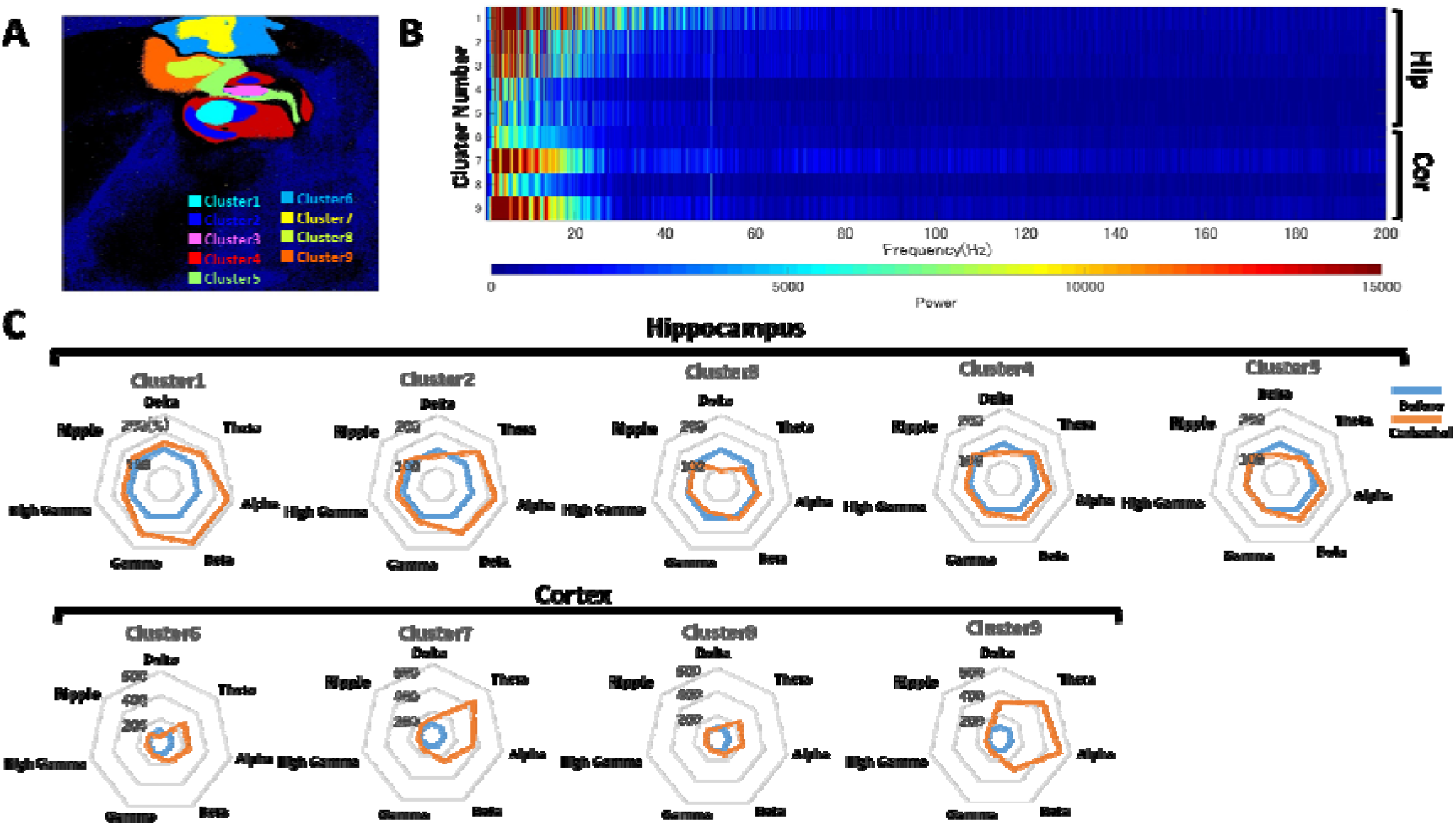
Region-Specific Frequency Characteristics Following Carbachol Treatment. (A) Cluster mapof the brain slice generated based on waveform similarity using k-means clustering and the elbow method. Each color-coded region corresponds to a specific cluster, reflecting anatomical locations. (B) Cluster-wise average frequency spectrum heatmap. Power spectra were calculated for each electrode within a cluster and then averaged to reveal frequency distribution across clusters. (C) Radar charts displaying frequency band power for each cluster. Values are normalized to the pre-treatment baseline (100%) for each cluster, allowing comparison of spectral changes induced by carbachol treatment.

## Result

### Field Potential Imaging of Acute Brain Slices Using an Ultra-high-density (UHD) CMOS-MEA

The UHD-CMOS-MEA, equipped with 236,880 electrodes spanning a 5.5 mm × 5.9 mm sensing area, enables large-scale recordings with both high spatial and temporal resolution. The complete recording setup, including the perfusion system, is illustrated in Figure 1A(a). Each electrode measures 10.52 µm × 10.52 µm, with an inter-electrode pitch of 1.2 µm, resulting in 81.3% coverage of the sensing surface (Figure 1A(b)).

Using this system, we recorded extracellular activity from acute sagittal mouse brain slices. The sensing area was sufficiently large to capture neural signals not only from the hippocampus and neocortex, but also from deeper regions such as the striatum, thalamus, midbrain, and substantia nigra, enabling simultaneous field potential imaging across nearly the entire brain slice (Figure 1A(c)). Figure 1B(a) displays a 30-second raster plot of spontaneous activity in the hippocampus and neocortex. A black band at 166 seconds, detected across 4,206 electrodes, indicates a hippocampal activation event, whereas a second band at 183 seconds, involving 8,860 electrodes, represents cortical activation. Figure 1B(b) shows potential heatmaps depicting the spatiotemporal propagation of extracellular activity across the hippocampal and cortical regions. Red and blue denote positive and negative potentials, respectively, corresponding to source and sink components of local field potentials. The anatomical structure of the hippocampus was clearly discernible from these potential maps. In the hippocampus, activity propagated from CA3 to CA2 and CA1 (Supplementary Movie 1(A,a)), with delayed peaks in positive potential observed across the circuit (Figure 1B(b), ➀–➆). During this propagation, positive potentials were predominantly detected in the pyramidal layer of CA3 (Figure 1B(c), ➀–➂) and the molecular layer of CA1 (Figure 1B(c), ➃–➆), whereas negative potentials were observed in the radiatum of CA3 (Figure 1B(c), ➇–➉) and the oriens of CA1 (Figure 1B(c), ⑭–⑯). Subsequently, the signal propagated to the dentate gyrus (DG) (Figure 1B(c), ⑪–⑬). In the neocortex, activation was initiated in the posterior parietal association areas (PTLp) (Figure 1B(c), ⑰).In the visual cortex (VIS), positive potentials were observed (Figure 1B(c), ⑱–◼ ), whereas in the somatosensory cortex (SS), negative potentials peaked during activation (Figure 1B(c), ◼–◼).

The UHD-CMOS-MEA yielded field potentials with a high signal-to-noise (S/N) ratio from both hippocampal and cortical regions, allowing for precise identification of sink–source relationships, particularly between CA3 and CA1.This system enables simultaneous mapping of spatiotemporal activity patterns in the hippocampus and neocortex, facilitating direct comparison with anatomical structures.

### Carbachol-Induced Firing Patterns of Beta Rhythm in the Hippocampal Region

Following the recording of spontaneous activity, carbachol treatment was applied to induce beta oscillations. Figure 2A(a) presents a 30-second raster plot of activity in the hippocampus and neocortex. A total of 41 oscillatory events were observed in the hippocampus, and 3 events in the cortex. In the hippocampus, carbachol treatment induced periodic oscillations at a frequency of 1.33□±□0.33 Hz. Figure 2A(b) shows raw extracellular waveforms recorded from the CA3 pyramidal layer between 8.5 and 13.5 seconds after treatment, revealing repetitive and periodic potential patterns. Figure 2A(c) illustrates the temporal evolution of potential maps during three representative oscillatory events (➃, ➆, ➇) from panel 2A(b). Although these events occurred at different time points, they exhibited a consistent propagation pattern: positive potentials originated in the CA3 oriens layer, followed by negative potentials in the radiatum and molecular layers. This activity subsequently propagated to the CA1 region, where negative potentials emerged in the CA3 pyramidal layer and extended into CA1. These propagation patterns were reproducible with millisecond precision across multiple events (Figure 2A(c), Supplementary Movie 2). These findings suggest that carbachol treatment reliably induced reproducible firing patterns within the hippocampus (Figure 2A(c)).

To further characterize these propagation patterns, we identified three distinct laminar pathways: from the CA3 pyramidal layer to the CA1 pyramidal layer (○), from the CA3 oriens layer to the CA1 radiatum (×), and from the CA3 molecular layer to the CA1 molecular layer (△) (Figure 2B). Figures 2C and 2D present representative raw waveforms for each pathway before and after carbachol treatment. Prior to treatment, reverberating activity was observed during oscillations in pathways ○ and ×. After treatment, the duration of oscillatory events was consistently reduced across all pathways, including △. The mean duration of oscillatory events post-treatment was 52.82□±□6.44 ms, corresponding to a frequency of 18.93□±□3.03 Hz, which falls within the beta frequency range.

Collectively, these results demonstrate that UHD-CMOS-MEA recording enables precise detection of repeated CA3-to-CA1 propagation patterns induced by carbachol treatment. The event durations were consistent with beta-frequency oscillations.

### Frequency Characteristics of the Hippocampus and Cortex Following Carbachol Treatment

To examine region-specific frequency characteristics in the hippocampus and cortex, we performed frequency analysis on a 5-second window, as indicated by the scale bar in Figure 2A(b). First, to identify spatial clusters of electrodes exhibiting similar waveform profiles, we applied k-means clustering to the raw extracellular waveforms recorded during the 5-second window. Figure 3A shows the resulting cluster map, where each electrode is color-coded according to waveform similarity.

Comparison with anatomical references (Allen Institute for Brain Science, 2015) revealed that the hippocampal region was subdivided into five clusters (Clusters 1–5) with clear laminar specificity: Cluster 1 corresponded to CA3 radiatum; Cluster 2 to CA3 pyramidal layer, stratum lacunosum-moleculare (Slm), and CA1 pyramidal layer; Cluster 3 to CA1 radiatum; Cluster 4 to CA3 oriens, dentate gyrus (DG), CA1 oriens, CA1 Slm, and subiculum pyramidal layer; and Cluster 5 to the CA2 region. Clusters 6–9 corresponded to cortical subdivisions as follows: visual cortex (Cluster 6, VIS); posterior parietal cortex (Cluster 7, PTLp); and somatosensory cortex (Clusters 8 and 9, SS).

Subsequent frequency analysis was performed for each cluster. The CA3 region, a key origin of signal propagation in the hippocampus, was primarily represented by Clusters 1 and 2. These clusters exhibited elevated power in the delta (1–3 Hz), theta (4–8 Hz), and alpha (8–15 Hz) bands, and also showed stronger beta (15–30 Hz) and gamma (35–50 Hz) components compared to other regions (Figure 3B). Following carbachol treatment, an increase in beta- and alpha-band power was observed in all hippocampal clusters (Clusters 1–5) (Figure 3C). Specifically, beta-band power increased to 183.2%, 152.7%, 105.1%, 125.9%, and 129.7% in Clusters 1 through 5, respectively. In the cortical clusters, beta-band power increased to 181.3% (Cluster 6); 240.2% (Cluster 7); 148.2% (Cluster 8); and 291.4% (Cluster 9). These results indicate that carbachol treatment enhanced beta-band activity across both hippocampal and cortical regions, enabling detailed analysis of beta-frequency distributions by anatomical subdivision. Furthermore, waveform-based clustering successfully captured structural distinctions across hippocampal and cortical layers, demonstrating that waveform morphology reflects region-specific electrophysiological features. Collectively, these findings confirm that UHD-CMOS-MEA-based field potential imaging is a powerful tool for investigating the relationship between induced beta rhythms and region-specific frequency dynamics in hippocampal–cortical circuits. Waveform-based clustering also revealed correspondence with anatomical structures in both the hippocampus and neocortex.

## Discussion

Using an ultra-high-density (UHD) CMOS microelectrode array (MEA) consisting of 236,880 electrodes spaced at 1.20 µm across a 5.5 mm × 5.9 mm sensing area, we successfully captured large-scale extracellular activity across multiple brain regions in acute mouse brain slices [Fig. 1A(c)]. Thanks to its exceptionally high spatiotemporal resolution, this platform enabled not only the precise tracking of activity propagation through hippocampal and cortical circuits, but also the delineation of anatomical laminar structures based on the spatial organization of field potential sinks and sources [Fig. 1B(b)].

Following carbachol treatment, we observed rhythmic oscillatory activity in the hippocampus at 1.33□±□0.33 Hz (Fig. 2A). The duration of individual events decreased to 52.82□±□6.44 ms, corresponding to a frequency of 18.93□±□3.03 Hz, indicating the successful induction of beta-range rhythms. Previous studies have reported that carbachol-induced beta oscillations arise primarily in pyramidal neurons within the CA3 region (Shimono et al., 2000). Consistent with this, our recordings revealed repeated CA3-originating oscillations that exhibited highly reproducible propagation patterns [Fig. 2A(c)]. Notably, the CA3-to-CA1 propagation sequences were temporally precise at the millisecond level and displayed consistent waveform morphology across events. Such fine-grained temporal and spatial resolution was made possible by the UHD-CMOS-MEA, which uniquely enabled the observation of these highly stereotyped activity patterns.

The hippocampus is well known for its role in memory formation and spatial information processing, with the CA3-to-CA1 pyramidal pathway constituting a fundamental conduit in this circuitry (Spruston, 2008). Prior to carbachol treatment, reverberating, seizure-like oscillations were observed along this pathway (Fig. 3C). However, carbachol administration suppressed these events and stabilized neuronal excitability, leading to more consistent and orderly propagation dynamics (Fig. 3D). These findings are consistent with prior reports that carbachol induces sustained rhythmic activity in cortical slices—likely by modulating cholinergic tone and suppressing runaway excitability (Schmidt et al., 2013). Given that beta rhythms are strongly associated with arousal and attentional states, our results suggest that carbachol treatment induced a wakefulness-like electrophysiological condition in acute brain slices.

Clustering based on waveform morphology proved to be a powerful approach for linking extracellular activity patterns with underlying anatomical and circuit-level architecture [Fig. 3A]. Notably, Cluster 5 encompassed the entire CA2 region and extended into CA1 and adjacent cortical areas. CA2 is known to act as a critical hub that integrates cortical inputs and relays information to CA1, playing a proposed role as a gate for transient inputs into memory networks and supporting temporal and sequential coding (Jones and McHugh, 2011). This anatomical distribution—spanning CA2 and reaching toward cortical regions—is consistent with these functional roles, and may reflect underlying circuit-level organization. These findings highlight the potential of waveform-based clustering to capture region-specific network features. In the cortex, beta-band oscillations have been widely associated with cognitive processes and sensory prediction, particularly through transient phase coupling between the visual cortex (VIS) and posterior parietal areas, including the PTLp (Vinck et al., 2016;Lohani et al., 2022). Consistent with these findings, our data demonstrated that carbachol treatment robustly enhanced beta-band power in cortical clusters (Fig. 3C), especially in VIS and PTLp. This enhancement may reflect transient coordination of activity across functionally connected cortical regions, potentially resembling aspects of higher-order cognitive states even under ex vivo conditions. Furthermore, the simultaneous acquisition of hippocampal and cortical activity within the same slice presents a valuable platform for exploring hippocampo-cortical interactions. Previous in vivo studies have shown that the hippocampus not only communicates with limbic and olfactory-related areas such as the entorhinal cortex, but also exhibits beta-band phase coupling with primary sensory cortices, including the visual cortex (Vinck et al., 2016)). In agreement with this, our data revealed concurrent beta-band enhancement in both hippocampal and cortical clusters following carbachol treatment (Fig. 3C), suggesting functional coordination between these regions. These findings underscore the potential of frequency-based cluster analysis to provide insights into large-scale network dynamics linked to higher cognitive functions.

Building on these findings, we employed field potential imaging using an ultra-high-density (UHD) CMOS-MEA—providing wide spatial coverage and high spatiotemporal resolution—to induce and characterize beta rhythms in the hippocampus and cortex of acute brain slices following carbachol treatment. Repetitive and stereotyped firing patterns originating from the CA3 region were observed, and beta-band power was enhanced in both hippocampal and cortical areas. Frequency analysis based on waveform-defined clusters enabled the characterization of neural dynamics that reflect both anatomical features and physiological function. Together, carbachol treatment and this FPI platform represent a promising approach for evaluating compound effects on hippocampal and cortical circuits under arousal-like conditions.

## Acknowledgments

This study was supported by the grant of collaborative project with Sony semiconductor solutions Inc.

## Author contributions

I.S and H.T. designed experiments; H.T. conducted experiments; H.T. and N.M. analyzed the data; I.S., H.T. and N.M. wrote the manuscript; I.S., H.T. and N.M. investigation; I.S. supervised the project.

## Supplementary Information

**Supplementary Movie 1. Propagation of a single hippocampal oscillatory event from CA3 to CA2 and CA1**.

This movie visualizes a oscillatory event recorded from an acute hippocampal slice using the ultra-high-density (UHD) CMOS-MEA. The color-coded map displays extracellular potentials recorded across 236,880 electrodes during a single oscillation cycle. The activity originates in the CA3 region and propagates sequentially to CA2 and CA1, illustrating the laminar spread of field potentials in real time. The color scale represents voltage amplitude (µV) recorded at each electrode.

**Supplementary Movie 2. Spatiotemporal dynamics of a beta-range oscillatory event propagating from CA3 to CA1**.

This movie presents a high-resolution visualization of extracellular potentials recorded during oscillatory events in the hippocampus, captured using the ultra-high-density CMOS-MEA following pharmacological treatment with carbachol (10□µM). Positive potentials originate in the CA3 oriens layer, followed sequentially by negative potentials in the radiatum and molecular layers. The activity subsequently propagates to the CA1 region, where negative potentials emerge in the CA3 pyramidal layer and extend into CA1. These propagation patterns were consistently observed with millisecond-level precision across multiple events. The color scale reflects the voltage amplitude (µV) recorded at each electrode.

